# Integrative Multi-Omics Analysis Reveals Stress-Specific Molecular Architectures in Soybean under Drought and Rust Infection

**DOI:** 10.1101/2025.07.07.663534

**Authors:** Gustavo Husein, Fernanda R. Castro-Moretti, Melina Prado, Lilian Amorim, Paulo Mazzafera, Javier Canales, Claudia B. Monteiro-Vitorello

## Abstract

Asian soybean rust (ASR), caused by *Phakopsora pachyrhizi*, represents a major constraint to soybean cultivation, with yield losses approaching 90% in the absence of effective control strategies. When coupled with the increasing incidence of drought driven by climate change, the co-occurrence of these biotic and abiotic pressures imposes a complex challenge for crop resilience. In this study, we explored the molecular responses of soybean (*Glycine max*) to concurrent water limitation and ASR infection through an integrative analysis of transcriptomic and metabolomic datasets. To capture both linear and conditional relationships among molecular features, we employed Weighted Gene Co-expression Network Analysis (WGCNA) alongside Copula Graphical Models (CGMs). WGCNA identified 17 gene co-expression modules exhibiting significant correlations with 27 annotated metabolites. Among these, abscisic acid showed consistent associations with drought-responsive modules enriched in central metabolic pathways and transcription factors such as Dof and bHLH. In contrast, modules linked to fungal infection were correlated with dipeptides and D-galacturonic acid, implicating early defense signaling and cell wall remodeling. The CGM framework further revealed sparse, condition-specific networks of differentially expressed genes and metabolites directly associated with each stressor, including genes encoding a dirigent-like protein, pentatricopeptide repeat proteins, and a nucleoredoxin, as well as metabolites such as inosine, Epi-dihydrophaseic acid and 2-oxoadipic acid. Notably, no gene or metabolite was found to be directly responsive to both stresses, underscoring the modular and stress-specific architecture of soybean defense. Together, these results highlight a hierarchical regulatory structure and demonstrate the value of combining correlation-based and dependency-driven models to identify candidate targets for multi-stress resilience breeding.

## 1. Introduction

Soybean (*Glycine max* (L.) Merr.) is one of the most important crops due to its high protein and oil content, supporting diverse applications in food, feed, and biofuel industries (Rahman et al., 2023). Despite its agronomic importance, soybean productivity is increasingly threatened by the occurrence of biotic and abiotic stresses. Asian soybean rust (ASR), caused by the biotrophic fungus *Phakopsora pachyrhizi*, is the most destructive soybean disease worldwide, capable of causing yield losses between 20% and 90% in the absence of chemical control, which to date remains the only efficient method to control this disease as there are no resistant genotypes (Dean et al., 2012). Climate change is expected to exacerbate this threat by expanding the geographic distribution of pathogens and intensifying selection pressure on host resistance genes (Alves et al., 2011; Burdon and Zhan, 2020; Delgado-Baquerizo et al., 2020; Ghini et al., 2007). Simultaneously, water limitation stress is projected to increase in severity due to shifts in precipitation patterns and prolonged drought periods (Leng and Hall, 2019; Thornton et al., 2014). The combined occurrence of abiotic and biotic stresses not only intensifies physiological damage, such as photosynthetic inhibition and premature defoliation (Echeveste Da Rosa, 2015), but also leads to complex and poorly understood interactions at the molecular level. These stress combinations are predicted to become more frequent and severe, yet their joint effects on plant development, metabolism, and defense remain insufficiently characterized (Camejo et al., 2005; Gerós et al., 2016; Mittler, 2006; Trivedi et al., 2022). Understanding the genetic and molecular basis of soybean responses to these concurrent stresses has therefore become a critical objective for improving crop resilience under current and future climatic scenarios (Kakumanu et al., 2012; Le et al., 2012; Xue et al., 2013).

Traditional approaches to elucidating molecular plant stress responses have typically relied on single-omics, focusing independently on transcriptomic or metabolomic data. While valuable, these strategies offer a fragmented view of molecular regulation, failing to capture how transcriptional and metabolic responses are temporally and functionally coordinated. Multi-omics frameworks have emerged as a powerful alternative, offering the possibility to map integrative regulatory circuits that span multiple biological layers (Roychowdhury et al., 2023; Sarfraz et al., 2025). Among the available tools, weighted gene co-expression network analysis (WGCNA) is widely used to identify gene modules co-expressed under specific conditions and to relate them to phenotypic or metabolic traits (Eicher et al., 2020; Langfelder and Horvath, 2008; Sanches et al., 2024). However, as a correlation-based approach, WGCNA does not distinguish between direct and indirect associations, which may limit its capacity to resolve causally relevant components (Kao et al., 2025; Ovens et al., 2021). Probabilistic graphical models represent an alternative strategy, providing a principled framework to infer conditional dependencies, which are associations that remain after accounting for the influence of all other measured variables (Behrouzi et al., 2020a; Farnoudkia and Purutcuoglu, 2019).

Among these strategies, Copula Graphical Models (CGMs) extend this concept to non-Gaussian and potentially non-linear dependencies, making them well suited to transcriptomic and metabolomic datasets that often violate assumptions of normality (Dobra and Lenkoski, 2011). By estimating sparse precision matrices in a copula-transformed space, CGMs enable the identification of direct relationships that remain after accounting for all marginal dependencies, thereby revealing a network of true conditionally independent associations among genes, metabolites, and experimental factors (Liu et al., 2009). Beyond their theoretical appeal, CGMs also offer practical advantages: they can handle missing data without the need for imputation, support the joint modeling of heterogeneous variable types, including both continuous and categorical features, and enable an exploratory, data-driven investigation of complex molecular systems (Behrouzi et al., 2020b; Behrouzi and Wit, 2019; Hermes et al., 2024). This latter aspect is particularly relevant in multi-omics studies, as it mitigates biases introduced by prior variable selection and allows previously unanticipated interactions to emerge from the structure of the data itself. Despite their potential, CGMs remain underutilized in plant systems biology. While recent studies have applied CGMs to assess the impact of drought stress on maize and wheat yield (Hermes et al., 2023), to dissect disease resistance in an interspecific raspberry population (Prado et al., 2024), and to identify the direct associations between dietary and epigenetic ageing in humans (Grootswagers et al., 2024a), their use in multi-omics integration, particularly under combined biotic and abiotic stress conditions in plants, remains virtually unexplored.

Building on previous single-omic studies of soybean responses to water limitation and *P. pachyrhizi* infection (Castro-Moretti et al., 2024; Husein et al., 2025), the present study expands upon these finds by applying an integrative multi-omics strategy that combines transcriptomic and metabolomic profiling under biotic and abiotic stress conditions. Using both WGCNA and CGM, we investigate whether the plant’s molecular response architecture is modular, stress-specific, or convergent under simultaneous stress exposure. This dual-framework approach enables the delineation of co-regulated gene–metabolite modules, the identification of direct associations relevant to each treatment condition, and the prioritization of candidate molecular components for future functional validation. Our findings provide deeper insights into the modular and hierarchical nature of plant stress responses and offer potential molecular targets for developing soybean cultivars with enhanced resilience to multiple environmental stresses.

## 2. Materials and methods

### 2.1. Experimental Design and Data Generation for Integrative Analysis

The transcriptomic and metabolomic data used in this study were generated from a single, comprehensive experiment for which the detailed methodology has been described in previous publications (Castro-Moretti et al., 2024; Husein et al., 2025). Briefly, the experiment followed a fully randomized factorial design with three biological replicates per treatment, using the susceptible soybean cultivar BMX Lança IPRO. Plants were grown in a greenhouse until the V4 developmental stage under two water availability regimes: a moderate water deficit (65% of the plant-available soil water) and a well-watered control (80%). Water treatments were initiated 48 hours prior to fungal inoculation. Urediniospores of *Phakopsora pachyrhizi* (10^5 spores mL^-1^) were sprayed onto the foliage while mock plants received sterile surfactant. Only leaf material collected at 12 and 24 hours after inoculation (HAI) from each treatment combination (water status × inoculation), which represent the time points common to both the metabolomic and transcriptomic datasets, were included in the present integrative analysis.

Untargeted metabolomic profiling was performed on V4 leaves using liquid chromatography coupled to mass spectrometry (LC–MS), as detailed by Castro-Moretti et al. (2024), enabling the simultaneous detection of primary and secondary metabolites. Transcriptomic data were generated from V3 leaves using RNA sequencing of poly-A-enriched libraries on the Illumina platform, following the procedures described in Husein et al. (2025), with gene-level quantification obtained through read alignment and transcript assembly.

### 2.2. Inference of Gene–Metabolite Associations via Linear Correlation

Raw gene counts were normalized for library size using the size factor method implemented in DESeq2 (Love et al., 2014). Outlier samples were identified by hierarchical clustering (hclust, method = “average”), based on sample-to-sample distances, resulting in the exclusion of two samples and a final dataset of 22 samples used for WGCNA construction. Genes were filtered to remove those with low expression (minimum expression count ≥ 1 in at least 25% of samples) and low variance (retaining the 75% most variant genes).

The WGCNA R package was used to construct a signed hybrid co-expression network and to calculate module eigengenes (MEs). Parameters included: soft-thresholding power (β) = 8, selected based on scale-free topology fitting; maxBlockSize = 44,000; networkType = ‘signed hybrid’; correlation type = ‘pearson’; and mergeCutHeight = 0.15, empirically chosen to yield a biologically interpretable number of modules, merging eigengenes with pairwise correlation above 0.85. Hub genes were identified using module membership (MM), representing the Pearson correlation between a gene’s expression and the corresponding ME.

Linear correlation analysis (Pearson correlation) was performed between MEs and the abundance profiles of the annotated metabolites using the corPvalueStudent function from the WGCNA package. Significant correlations were determined using a strict threshold (p < 0.0001), chosen to ensure conservative selection under high-dimensional correlation testing. Significant metabolites were then subjected to hierarchical clustering (hclust, method = ‘average’) applied to the transposed matrix of significant ME-metabolite correlations, using a distance cut-off (h = 1.5) to group metabolites based on their correlation patterns across modules. This threshold was chosen empirically to yield a manageable number of biologically interpretable clusters for downstream analysis.

### 2.3. Inference of Gene–Metabolite Associations via Copula Graphical Models

To explore conditional dependencies, a Copula Graphical Models (CGM) approach was applied using the nutriNetwork R package (Behrouzi, 2023). Given the high computational demand of estimating high-dimensional precision matrices, we restricted the input gene set to the unique differentially expressed genes (DEGs) identified across any of the three experimental contrasts (Fungi, Water and Interaction) and both time points, using DESeq2 (|log_2_FC| > 1, adjusted p-value < 0.05, Benjamini–Hochberg correction). The same pre-processed expression matrix used in the linear correlation analysis (Section 2.2), with outlier samples removed and low-expression genes filtered, was used as the basis for DEG selection.

The input dataset included: (i) expression profiles of the 657 DEGs identified, (ii) abundance profiles of the 455 annotated metabolites, and (iii) a binary incidence matrix encoding the experimental treatment conditions (Fungal Inoculation: 0=No, 1=Yes; Water Limitation: 0=No, 1=Yes; Sampling Time: 0=12 HAI, 1=24 HAI). This resulted in a 1,115-variable dataset across the 22 samples. All variables (gene expression, metabolite abundance, incidence matrix columns) were standardized to zero mean and unit variance prior to network construction.

The nutriNetwork function was executed using the Gibbs sampling algorithm (method = “gibbs”) with a fixed regularization parameter (rho = 0.3), selected to balance sparsity and biological interpretability. Other parameters included em.iter = 5 and em.tol = 0.001. This procedure estimates the copula-based precision matrix (inverse covariance matrix), where non-zero off-diagonal elements represent partial correlations (conditional dependencies) between variables after conditioning on all others. Unlike correlation-based approaches, this allows for the identification of direct associations, removing indirect effects mediated by other variables and providing a more accurate representation of the underlying regulatory structure. The resulting model was post-processed using the selectnet function to extract the final adjacency matrix representing the inferred conditional dependence network. Edges directly connecting the ‘Fungal Inoculation’ or ‘Water Limitation’ nodes to genes or metabolites were extracted to identify variables directly associated with each stress condition within the network structure.

### 2.4. Functional Enrichment Analysis

KEGG pathway enrichment analysis was performed using the SoybeanGDB enrichment tool (https://venyao.xyz/SoybeanGDB/, accessed March 28, 2025), as described by Li et al. (2023). Annotation of soybean transcription factors (TFs) was obtained from PlantTFDB 4.0 (Glycine max TF list, downloaded from https://planttfdb.gao-lab.org on March 28, 2025) (Jin et al., 2017), following a characterization of the top 10 hub genes per module. Resistance Gene Analog (RGA) classification and enrichment analysis were conducted following the methodology described by Rody et al. (2019), calculating PTI and ETI enrichment scores relative to the total number of RGAs per module and in the genome. General gene functional annotation was based on the Wm82.a4.v1 genome annotation from JGI Phytozome portal (Glycine max Wm82.a4.v1, accessed on March 28, 2025) (Goodstein et al., 2012).

## 3. Results

The integration of transcriptomic and metabolomic data using two distinct methodologies enabled the identification of stress-responsive gene-metabolite associations in soybean. The first approach, based on linear correlation analysis, identified marginal associations between co-expressed gene modules and metabolites, providing insight into coordinated metabolic and transcriptional regulation. The second approach, employing copula graphical models, detected probabilistic dependencies, capturing non-linear direct relationships between differentially expressed genes (DEGs) and metabolites under biotic and abiotic stress conditions.

### 3.1. Correlation-based dependency

A co-expression network was constructed from 29,346 genes expressed across two time points and multiple stress conditions using Weighted Gene Co-expression Network Analysis (WGCNA). After merging highly correlated modules (Pearson correlation > 0.85), 32 gene modules were obtained. Pearson correlations were then calculated between these gene modules and the 455 identified metabolites, using a strict significance threshold (p < 0.0001). This analysis identified 17 modules with significant associations to 27 metabolites (Figure 1).

**Figure 1.**
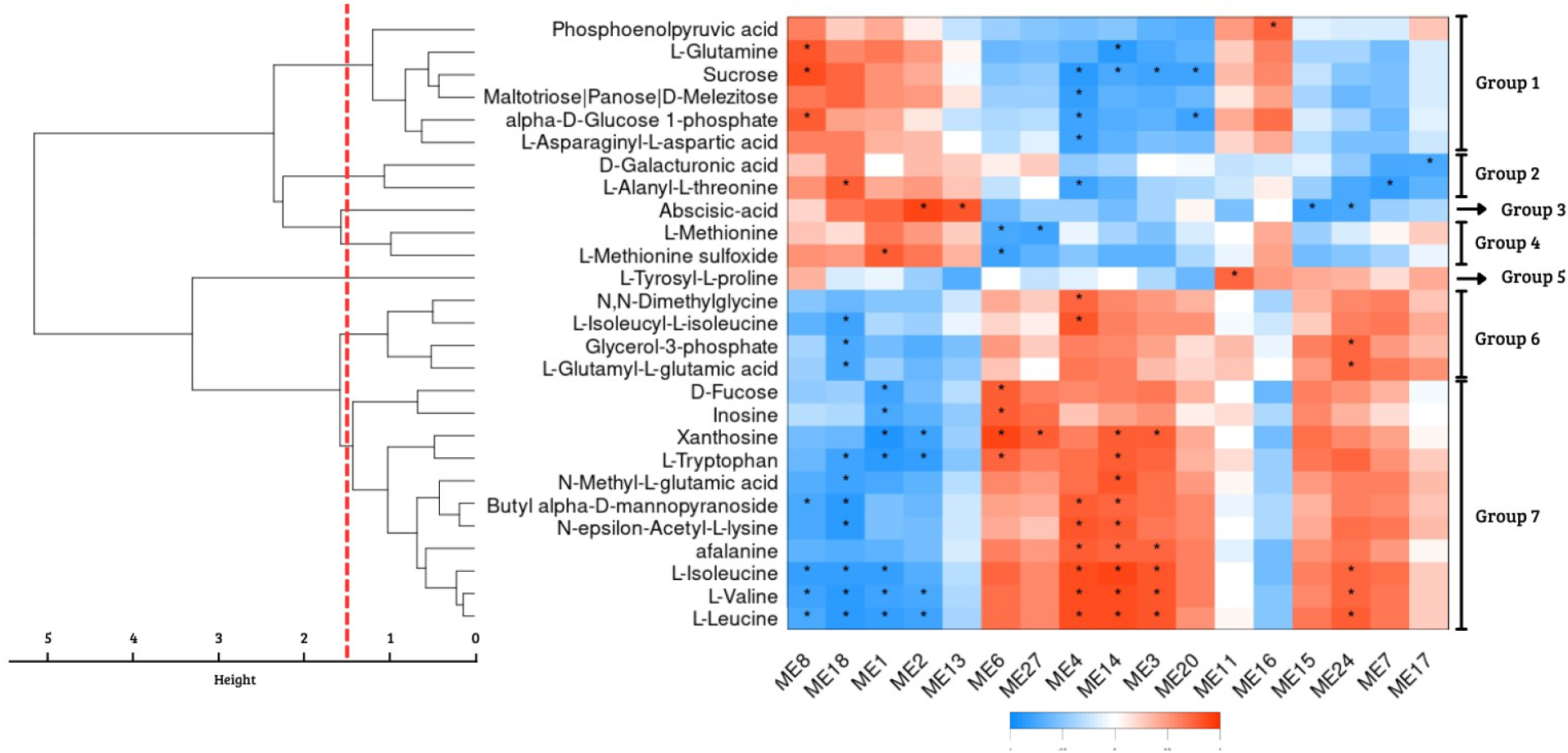
Significant associations between WGCNA modules and metabolites. Significant (p < 0.0001) linear association (Pearson correlations) of seventeen WGCNA modules with 27 annotated metabolites. The heatmap displays scaled correlation values between module eigengenes (MEs) and metabolite abundances, with positive correlations shown in red and negative correlations in blue. Asterisks (*) indicate statistically significant associations. Hierarchical clustering of metabolites was performed using complete linkage and Euclidean distance, and the dendrogram was cut at a height of 1.5, defining seven distinct metabolite clusters (Groups 1 to 7), labeled on the right.

These significant metabolites were clustered into seven groups based on hierarchical clustering with a distance threshold of 1.5 (Figure 1), Group 1 containing six metabolites (Phosphoenolpyruvic acid, L-glutamine, Sucrose, Maltotriose/Panose/D-melezitose, alpha-D-glucose 1-phosphate and L-asparaginyl-L-aspartic acid), Group 2 with two (D-galacturonic acid and L-alanyl-L-threonine), Group 3 with one (Abscisic acid), Group 4 with two (L-methionine and L-methionine sulfoxide), Group 5 with one (L-tyrosyl-L-proline), Group 6 with four (N,N-dimethylglycine, L-isoleucyl-L-isoleucine, Glycerol-3-phosphate and L-glutamyl-L-glutamic acid) and Group 7 with eleven (D-fucose, Inosine, Xanthosine, L-tryptophan, N-methyl-L-glutamic acid, Butyl alpha-D-mannopyranoside, N-epsilon-acetyl-L-lysine, Afalanine, L-isoleucine, L-valine and L-leucine).

From the identified metabolite clusters, three emerged as particularly relevant based on their linear associations with transcriptomic modules (**Figure 2**): group 3 was classified as responses to abiotic stress, whereas groups 2 and 5 were associated with biotic stress response.

**Figure 2.**
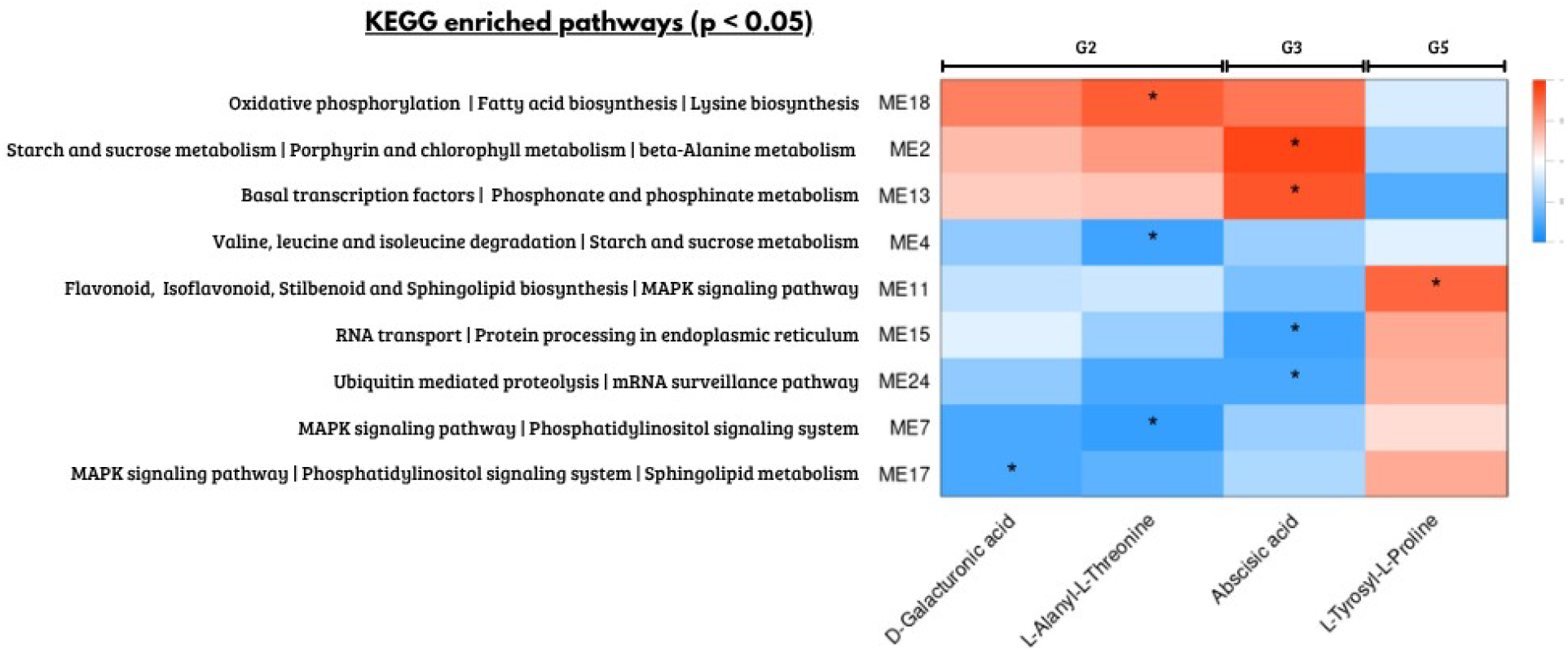
KEGG enriched pathways of gene modules significantly correlated with stress-associated metabolites. Heatmap showing summary KEGG pathways (adjusted p < 0.05) enriched in each module with significant linear correlations (Pearson, p < 0.0001) between annotated metabolites from Groups 2, 3 and 5. Metabolites were grouped based on correlation patterns, and significant correlations are indicated by asterisks. Red indicates positive correlations and blue negative correlations.

To further explore the biological roles of the modules associated with stress-responsive metabolites, we assessed their enrichment in resistance gene analogs (RGAs) and characterized the transcription factor (TF) families represented among their hub genes (Table 1, Supplementary Table 1).

**Table 1.** Functional characterization of gene co-expression modules significantly associated with stress-responsive metabolites.

| Modules | TF Class | PTI enrichment | ETI enrichment |
| --- | --- | --- | --- |
| ME18 | — | 5.10 | 7.00 |
| ME2 | — | — | — |
| ME13 | Dof, bHLH | — | — |
| ME4 | FAR1 | — | — |
| ME11 | — | — | — |
| ME15 | — | 3.53 | 4.96 |
| ME24 | gATA | 1.76 | — |
| ME7 | WRKY | — | — |
| ME17 | NAC, MYB, CAMTA | — | — |

Resistance gene analog (RGA) enrichment scores for selected modules were calculated as fold enrichment of genes involved in pattern-triggered immunity (PTI) or effector-triggered immunity (ETI), based on their relative frequency compared to genome-wide RGA distribution. Transcription factor (TF) families were identified
among the top 10 hub genes within each module using known TF class annotations. A dash (—) indicates the absence of TFs or no significant RGA enrichment.

Group 3, represented by Abscisic acid, showed a significant linear association with four gene co-expression modules. Modules 2 (r = 0.893) and 13 (r = 0.805), containing 2,556 and 642 genes respectively, were positively correlated. Their KEGG enrichment (Supplementary Table 2) included pathways such as starch/sucrose metabolism, porphyrin and chlorophyll biosynthesis. Conversely, modules 15 (r = -0.767) and 24 (r = -0.734), comprising 539 and 207 genes, showed negative correlations and were enriched in processes such as RNA transport, protein processing and ubiquitin-mediated proteolysis. These modules were also enriched for resistance gene analogs (RGAs) related to pattern-triggered immunity (PTI) and effector-triggered immunity (ETI). In terms of transcription factors, DNA binding with one finger (Dof) and basic helix-loop-helix (Bhlh) families were enriched in the positively associated modules, while the GATA family was predominant among the negatively associated ones. At the metabolite level, ABA showed increased abundance under water-limiting conditions and reduced abundance in inoculated plants, when compared with the control treatment, exclusively at 12 HAI, with no evident differences between treatments at 24 HAI (Figure 3).

**Figure 3.**
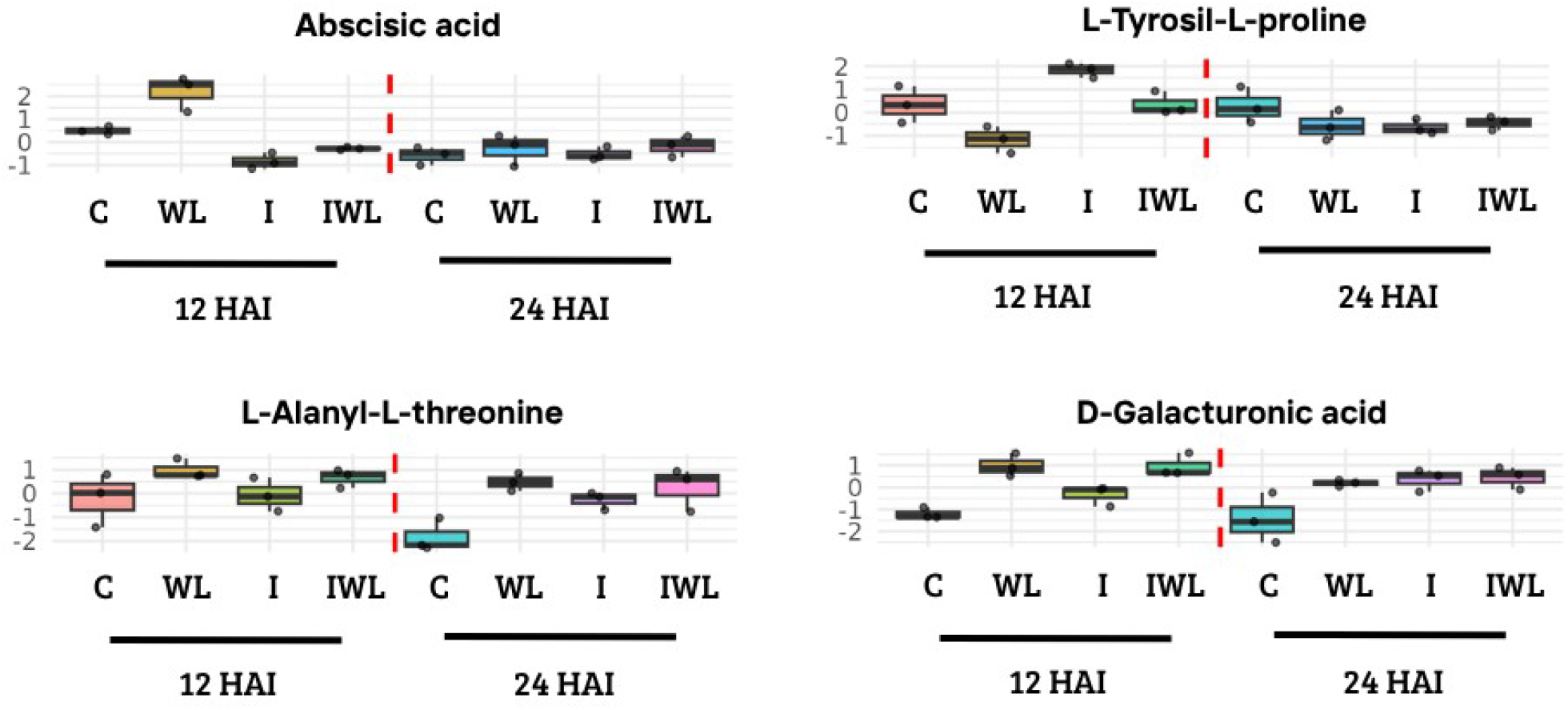
Abundance profiles of selected metabolites across treatments and time points. Boxplots showing the normalized abundance of four metabolites (Abscisic acid, L-tyrosyl-L-proline, L-alanyl-L-threonine, and D-galacturonic acid) in soybean leaves under different treatments: control (C), water limitation (WL), fungal infection (I), and combined stress (IWL), evaluated at 12 and 24 hours after inoculation (HAI). Colors represent treatment groups, and each dot corresponds to an individual biological replicate. Metabolites were selected based on their stress-specific associations and correlations with gene co-expression modules.

Group 5, composed of the metabolite L-tyrosyl-L-proline, was only significantly positively correlated with module 11 (r = 0.754), which contains 2,235 genes. This module was enriched in KEGG pathways (Supplementary Table 2) related to flavonoid, isoflavonoid, and stilbenoid biosynthesis, as well as sphingolipid metabolism and MAPK signaling pathway, while no enrichment of RGAs or transcription factor families was observed. Consistent with this association, L-tyrosyl-L-proline showed higher abundance in inoculated plants at 12 hours after inoculation (HAI) and reduced abundance under water-limiting conditions. At 24 HAI, no clear differences in abundance were observed between treatments (Figure 3).

Finally, D-galacturonic acid and L-alanyl-L-threonine, components of Group 2, were positively correlated with module 18 (r = 0.604; r = 0.774), containing 407 genes, and negatively correlated with modules 4 (r = -0.463; r = -0.766; 2,235 genes), 7 (r = -0.721; r = -0.827; 1,381 genes) and 17 (r = -0.755; r = -0.647; 434 genes). Positively correlated KEGG pathway enrichment (Supplementary Table 2) included oxidative phosphorylation, fatty acid biosynthesis, and lysine biosynthesis, while negatively correlated modules were enriched in pathways associated with branched-chain amino acid degradation, autophagy, starch and sucrose metabolism, and MAPK signaling. The positively associated module also showed significant enrichment for RGAs involved in PTI and ETI responses, while no RGA enrichment was found for negatively correlated modules. Among transcription factors, only negatively associated modules were enriched in TF of families FAR1, WRKY, CAMTA, NAC, and MYB. At the metabolite level, L-alanyl-L-threonine exhibited increased abundance in both fungal and water-limited treatments at 24 HAI, with no marked differences at 12 HAI. D-Galacturonic acid showed consistently higher abundance under both biotic and abiotic stress conditions at both 12 and 24 HAI (Figure 3).

### 3.2. Copula graphical models

A second analytical strategy was employed to detect non-linear and conditional dependencies between genes and metabolites using a copula graphical model. Due to the computational intensity of this method, gene expression data were first filtered to include only differentially expressed genes (DEGs).

A total of 703 DEGs (|log_2_FoldChange| > 1, padj < 0.05) were identified across the two collection time points (12 and 24 HAI) and three experimental contrasts: fungal infection by *Phakopsora pachyrhizi*, water limitation, and the interaction of both stresses. Most DEGs were condition-specific, while a subset was shared across two or more contrasts and/or time points (Figure 4). Among these, 545 DEGs were associated with the biotic (Inoculated vs. non-inoculated), 141 with abiotic stress (water-limited vs. control), and 17 with the interaction between the two stress types. When stratified by sampling time, the majority of DEGs were observed at 12 HAI (582 genes, 82.78%), with a smaller subset detected at 24 HAI (121 genes, 17.21%) (Supplementary Table 3).

**Figure 4.**
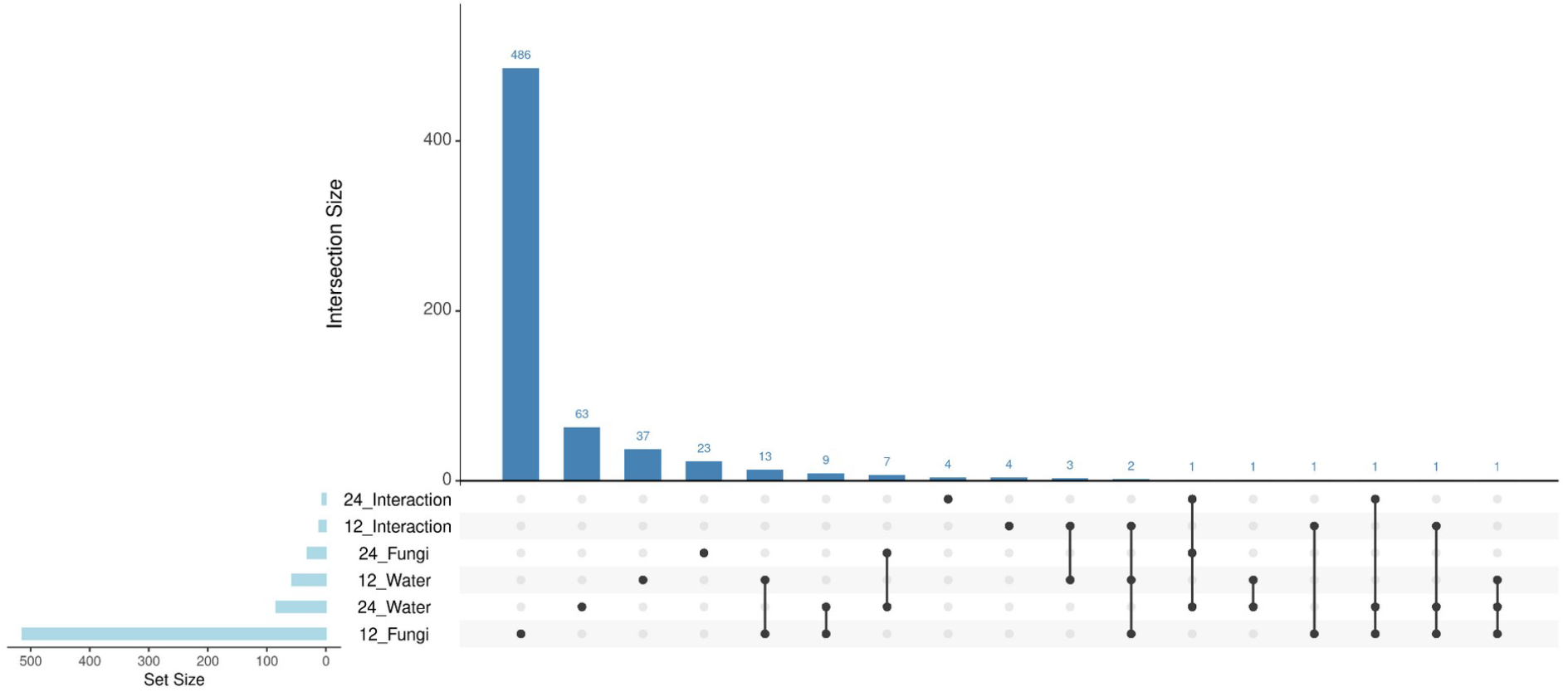
Intersection of differentially expressed genes (DEGs) across stress contrasts and time points. UpSet plot showing the number of DEGs identified under each condition, defined by three contrasts (Fungi infection, Water limitation, Interaction of both stresses) and two time points (12 and 24 HAI). Each horizontal bar on the left represents the total number of DEGs in each condition. The matrix below the vertical bars indicates the combinations of conditions, where shared filled circles denote the presence of an intersection. Vertical bars represent the number of DEGs shared across the respective combination of conditions.

The 657 unique DEGs were integrated with the 455 annotated metabolites and the binary incidence matrix (indicating stress condition, abiotic and biotic, and time) to construct a 1115-variable dataset. The copula graphical model, applied to this matrix, identified partial correlations among variables, revealing conditional dependency structures beyond simple linear associations. Variables were filtered to retain only those directly correlated with the biotic (FUNGI) and abiotic (WATER) stress conditions (Figure 5A, Supplementary Table 4).

**Figure 5.**
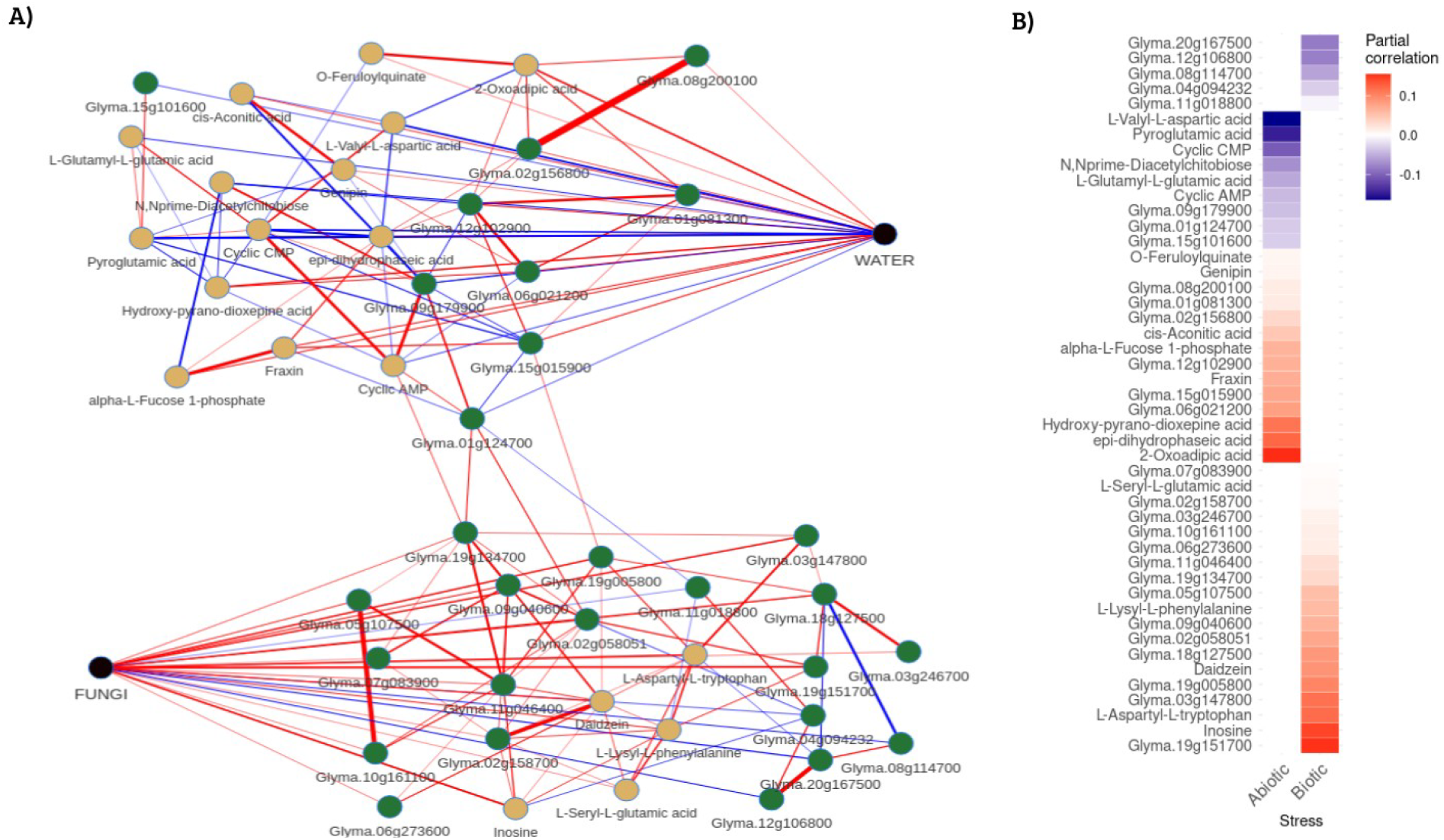
Conditional dependence network inferred by copula graphical modeling. (A) Subnetwork extracted from the full copula graphical model showing only the variables directly connected to the biotic (FUNGI) or abiotic (WATER) stress nodes. Nodes represent genes (green) and metabolites (yellow), color-coded according to the direction of their partial correlation with the respective stress condition (blue: negative; red: positive). The central nodes labeled FUNGI and WATER correspond to the binary incidence variables included in the model. Only direct (first-order) associations are displayed. (B) Heatmap displaying the magnitude and direction of partial correlations (ρ) for each gene and metabolite directly connected to either the Abiotic (left) or Biotic (right) condition. Positive correlations are shown in red, and negative correlations in blue. Variables are grouped by stress condition, with genes and metabolites ordered by correlation strength within each group.

From the copula-based network, a total of 19 genes, 14 with positive and 5 with negative partial correlations, and 5 positively associated metabolites were identified as being connected to fungal stress (Supplementary Table 4). For the water deficit condition, 9 genes (6 positive, 3 negative) and 14 metabolites (8 positive, 6 negative) were identified (Supplementary Table 4). Across both stress conditions, partial correlation coefficients ranged from 0.155 to –0.164 (Figure 5B, Supplementary Table 4). Notably, no gene or metabolite was found to be directly associated with both stresses

For fungal infection, the highest positive correlations were found for the genes *Glyma*.*19g151700* (ρ = 0.154, Pentatricopeptide repeat superfamily protein) and *Glyma*.*03g147800* (ρ = 0.117, Disease resistance-responsive (dirigent-like protein) family protein), and for the metabolite Inosine (ρ = 0.145). The most negative correlation was observed with the gene *Glyma*.*20g167500* (ρ = –0.084, Abscisic stress ripening-like protein).

In response to water stress, the highest positive correlations were observed for the metabolite 2-Oxoadipic acid (ρ = 0.155), Epi-dihydrophaseic acid (ρ = 0.124), Hydroxy-pyrano-dioxepine acid (ρ = 0.114) and the gene *Glyma*.*06g021200* (ρ = 0.081, Probable nucleoredoxin 2-like isoform 2). The most negative partial correlations were observed for L-Valyl-L-aspartic acid (ρ = –0.164), Pyroglutamic acid (ρ = –0.147), and for the gene *Glyma*.*09g179900* (ρ = –0.040, Receptor-like kinase).

Although each integrative strategy identified largely distinct gene–metabolite relationships, a few overlapping features emerged, reinforcing the biological consistency of the integrative findings. The gene *Glyma*.*08g114700* (calmodulin-binding protein) exhibited a negative partial correlation with the presence of the fungus in the CGM (ρ = –0.058), and was also a member of WGCNA Module 17, which was negatively correlated with Group 2 (metabolite cluster enriched in biotic response signatures). Similarly, *Glyma*.*06g021200* (probable nucleoredoxin), *Glyma*.*12g102900* (beta-amylase 5), *Glyma*.*01g081300* (alpha-glucosidase), and *Glyma*.*08g200100* (HAD superfamily phosphatase) were positively associated with the water-limiting condition in the CGM and simultaneously located in WGCNA modules positively correlated with Group 3, represented by abscisic acid.

## 4. Discussion

The integrative strategy employed in this study, combining transcriptomic and metabolomic data based on linear associations (Pearson correlation) and conditional dependency inference (CGM), revealed discrete regulatory architectures underlying soybean responses to biotic and abiotic stress, caused by ASR and water limitation, respectively. Notably, no gene or metabolite was directly associated with both stress conditions, supporting a model in which soybean activates stress-specific modules rather than shared signaling pathways under concurrent challenges. This separation was consistently observed in both the correlation-based and conditional models, suggesting that each stressor elicits a dedicated molecular response with minimal overlap. Interestingly, the response uncovered by the integrated analysis differed from those observed in the transcriptomic and metabolomic datasets individually: while the transcriptome revealed a unique expression state under stress combination (Husein et al., 2025), and the metabolome showed time-dependent modulations of specific compounds (Castro-Moretti et al., 2024), the current multi-omic approach, in contrast, highlights the preservation of stress-specific responses even under simultaneous exposure, reflecting a modular regulatory strategy likely evolved to mitigate antagonistic crosstalk between signaling pathways such as those mediated by abscisic acid (ABA) versus pathogen-induced responses, optimizing resource allocation depending on the nature of the environmental signal (Anderson et al., 2004). The findings reinforce the need for integrative approaches that distinguish between coexpression and direct interaction, especially when investigating multifactorial stress contexts.

Among all metabolites detected, ABA was the most consistently associated with the water-limiting treatment, identified by both linear correlation and CGM approaches. This finding is in line with the central role of ABA in plant drought responses, as it is widely recognized as the primary phytohormone mediating adaptive responses to water deficit through rapid accumulation, stomatal closure, and transcriptional reprogramming, resulting on the expression of drought-responsive genes (Margay et al., 2024; Muhammad Aslam et al., 2022). In the correlation-based analysis, ABA was positively correlated with modules enriched in primary metabolic pathways, such as starch and sucrose metabolism, which are commonly affected under drought stress (Nidumolu et al., 2023). The presence of Dof and bHLH transcription factors (TF) in these modules suggests regulatory coordination of metabolic and developmental pathways under ABA influence. Prior studies have demonstrated that these TF families promote drought tolerance in tomato and peanut, primarily by enhancing ABA accumulation, increasing ROS scavenging, and repressing ABA catabolic activity (Liang et al., 2022; Liu et al., 2025). In contrast, modules showing negative correlations with ABA were associated with RNA and protein-related processes, including ubiquitin-dependent protein degradation. This negative association likely reflects a shift in regulatory priorities under drought-induced ABA signaling. As reviewed by Xu et al. (2024), elevated ABA levels modulate the ubiquitin–proteasome system (UPS) to promote the selective degradation of transcriptional regulators and signaling components, prioritizing stress adaptation over growth-related processes. These modules also exhibited enrichment for resistance gene analogs (RGAs) from both PTI and ETI classes. Although RGAs are typically implicated in biotic defense, their repression in response to water deficit suggests an antagonistic regulation between abiotic and biotic responses, possibly mediated by ABA-induced suppression of defense pathways. Supporting this view, ABA was shown to promote immune suppression in *Arabidopsis* through the induction of HAI-type phosphatases that inactivate immune-associated MAPKs, revealing a mechanistic basis for ABA-mediated antagonism of immune pathways (Mine et al., 2017).

The CGM analysis yielded a narrower and more stringent subset of directly associated features, identifying nine genes and fourteen metabolites with direct conditional dependencies on the water stress factor. Among these, Glyma.06g021200, encoding a nucleoredoxin, was directly linked to the water condition node and may act as a key modulator of redox homeostasis under water limitation. Nucleoredoxins are oxidoreductases known to preserve the activity of antioxidant enzymes, such as catalase under ROS-enriched conditions. In Arabidopsis, the activity of catalase under oxidative conditions is supported by NRX1, which helps preserve its functionality during oxidative stress (Kneeshaw et al., 2017). In wheat, overexpression of TaNRX1 enhances drought tolerance by boosting osmoprotectant accumulation, increasing antioxidant enzyme activity, and modulating stress-related gene expression (Zhang et al., 2021). These observations align with the hypothesis that Glyma.06g021200 may act independently of ABA signaling to maintain redox balance under water limitation conditions. Additionally, the presence of metabolites such as 2-oxoadipic acid and epi-dihydrophaseic acid (epi-DPA) suggests targeted metabolic regulation under water limitation conditions. The former, involved in lysine and tryptophan degradation, was also found in roots of drought-stressed pine seedlings, implying a contribution to amino acid turnover and redox buffering (Wu et al., 2023). The latter is a byproduct of ABA catabolism, accumulated during ABA deactivation, marking the cessation of ABA signaling and the transition to homeostatic recovery (Bai et al., 2022). Together, the presence of redox-related genes and hormone-catabolizing metabolites indicates a coordinated stress response involving both ABA deactivation and metabolic adjustment to restore redox equilibrium.

In contrast to the metabolite-dominated response observed under water limitation, the response to *P. pachyrhizi* infection was dominated by transcriptional reprogramming. The CGM analysis identified nineteen genes and five metabolites conditionally dependent on the biotic treatment, including genes encoding a disease resistance-responsive (dirigent-like protein) family and pentatricopeptide repeat (PPR) proteins. Dirigent-like proteins, such as CaDIR7 in *Capsicum annuum*, are involved in lignin biosynthesis and defense against *Phytophthora spp*., reinforcing cell walls to hinder pathogen invasion (Khan et al., 2018). Similarly, PPR proteins mediate organellar gene expression by controlling RNA metabolism, and several have been implicated in defense signaling and stress adaptation via chloroplast and mitochondrial regulation (Meng et al., 2024). The limited number of associated metabolites suggests that the early stages of biotic stress response for the susceptible soybean cultivar favor signal perception and transcriptional reprogramming over broad metabolic shifts, a pattern consistent with rapid activation of immune regulatory cascades (Meraj et al., 2020). Interestingly, modules negatively correlated with biotic stress-related metabolites identified using the correlation-based approach were enriched for WRKY, NAC, MYB, and CAMTA transcription factors, which are key regulators of biotic and abiotic stress response (Bian et al., 2020; Li et al., 2024), but lacked enrichment for classical RGAs. This configuration likely reflects pathogen-mediated suppression of host signaling, which has been explicitly described in the *Glycine max– Phakopsora pachyrhizi* pathosystem, where secreted effectors suppress basal defenses to enable effector-triggered susceptibility (ETS) in compatible interactions (Chicowski et al., 2023; Gupta et al., 2023). The lack of RGA enrichment during these early stages, together with the expression of stress-related transcription factors, is consistent with observations in other susceptible soybean genotypes, where a limited or incomplete immune activation has been associated with reduced NLR expression, suggesting that attenuated immune signaling contributes to disease susceptibility (Hao et al., 2023).

Among the metabolites associated with the rust presence, inosine exhibited the strongest positive partial correlation with infection, suggesting a potential role in early immune regulation. In plants, the accumulation of inosine di- and triphosphate, resulting from a deficiency in inosine triphosphate pyrophosphatase (ITPA), leads to the accumulation of salicylic acid, an immune response, and premature senescence, indication a signaling function in stress perception conserved in many organisms (Straube et al., 2023). Moreover, in animal models, inosine has been shown to fine-tune innate immunity by suppressing pro-inflammatory cytokines such as IL-1β while enhancing complement system activation, thus coordinating cellular and humoral immune responses during bacterial infection (Jiang et al., 2022). Although the functional role of inosine in soybean immunity remains to be elucidated, its accumulation under fungal infection may reflect a conserved metabolic strategy across kingdoms to modulate biotic defense responses.

In the correlation-based integrative analysis, three metabolites emerged as particularly relevant for *P. pachyrhizi* infection, combining statistical robustness with biological relevance. Two of them, L-alanyl-L-threonine and L-tyrosyl-L-proline, suggest functional links to amino acid-mediated defense mechanisms. The former, though not directly linked to plant immunity, may accumulate as part of the host’s early biochemical response to fungal invasion, given the role of dipeptides in defense metabolite biosynthesis and immune signaling (Parthasarathy et al., 2021). L-tyrosyl-L-proline combines two stress-related amino acids: tyrosine, which is central to antioxidant systems and the production of lignin and phytoalexins (Xu et al., 2020), and proline, known for its osmoprotective and redox-balancing effects under stress (Meena et al., 2019). Although the functional role of L-tyrosyl-L-proline in soybean immunity is uncharacterized, its positive correlation with a module enriched in MAPK signaling and flavonoid-related biosynthesis supports its possible involvement in defense regulation. D-Galacturonic acid, also highlighted, is a component of oligogalacturonides, which act as elicitors of immune responses (Ferrari et al., 2013; Savatin et al., 2014). Though less active than the full oligomer, its accumulation may reflect pectin degradation initiated during fungal entry. This is consistent with the known infection dynamics of *P. pachyrhizi*, which establishes haustoria within the first 24 hours (Gupta et al., 2023). Together, these metabolites may represent distinct biochemical facets of the biotic stress response, potentially linking early signaling, metabolic adjustment, and structural remodeling in the absence of classical immune activation.

The use of both WGCNA and CGM enabled a multi-layered interpretation of gene– metabolite associations by distinguishing broad coexpression patterns from direct conditional dependencies. WGCNA revealed coordinated transcriptional modules associated with stress-induced metabolic changes, providing an overview of systemic regulatory responses. However, such correlation-based analyses capture both direct and indirect relationships, potentially obscuring key regulatory nodes. In contrast, CGM estimated conditional dependencies among variables, identifying sparse, high-confidence networks in which only direct interactions are retained. Importantly, even near-zero partial correlations may capture meaningful associations in the copula-transformed space, since these values are calculated after correcting for all other variables in the dataset (Grootswagers et al., 2024b; Rossell and Zwiernik, 2021; Xia and Li, 2021). The non-overlapping sets of genes and metabolites identified by each approach emphasize their analytical complementarity, and their integration offers a more accurate prioritization of molecular targets for functional studies under complex stress conditions. Future studies may benefit from adopting this dual-framework strategy as a standard in multi-omics integration, particularly when aiming to disentangle context-specific regulatory circuits in plant stress biology.

The stress-specific molecular signatures identified in this study provide a valuable resource for breeding programs aiming to enhance tolerance to abiotic and biotic stresses simultaneously. The absence of shared genes or metabolites between stress conditions implies that dual stress tolerance cannot be efficiently obtained by selecting for markers that are simultaneously associated with both stresses. Instead, targeted stacking of stress-specific regulators, carefully chosen to avoid antagonistic cross-regulation, emerges as a more promising strategy. Among the conditionally associated genes identified by CGM, *Glyma*.*06g021200* (nucleoredoxin), *Glyma*.*19g151700* (PPR protein) and *Glyma*.*03g147800* (dirigent-like protein) represent particularly compelling candidates for functional characterization. These genes are directly associated with abiotic (first gene) and biotic (last two genes) stress responses, and their respective roles in redox homeostasis, RNA metabolism, signaling, and pathogen response, position them as putative hubs for engineering enhanced stress resilience. Similarly, metabolites such as ABA, 2-oxoadipic acid, and inosine, which exhibited stress-specific associations, may serve as reliable biochemical markers for phenotyping early-stage responses in field conditions. The observed conditional independence between responses also suggests that improvements in one trait may not incur penalties in the other, provided that antagonistic regulators like ABA are carefully managed. These insights offer a rational framework for designing molecular breeding strategies that maximize multi-stress resilience while minimizing trade-offs. However, the use of only the susceptible cultivar inherently limits the generality of our conclusions. Future experiments that include resistant genotypes will be essential to disentangle susceptibility-specific effects from universal stress responses. Moreover, our focus on early time points (12 and 24 HAI) may have overlooked later-emerging molecular interactions. This way, extending sampling to later stages could uncover additional layers of drought–pathogen interplay. Lastly, the functional roles proposed for specific genes and metabolites, although supported by published evidence, remain speculative until validated in soybeans. Therefore, they should be viewed as promising targets for directed functional assays rather than confirmed regulators.

By integrating transcriptomic and metabolomic data through both correlation-based and conditional dependency frameworks, this study delineates distinct regulatory architectures underlying soybean responses to abiotic and biotic stress. The identification of stress-specific modules, the central role of ABA in drought adaptation, and the transcriptional reprogramming observed during pathogen challenge highlight the modular and hierarchical nature of these responses. The complementarity between WGCNA and CGM proved essential for refining candidate genes and metabolites, offering a robust platform for future functional validation. These findings not only advance our understanding of how plants orchestrate specialized responses to complex environmental stimuli but also provide actionable molecular targets for developing multi-stress-resilient cultivars through integrative breeding and synthetic biology approaches.

## Acknowledgments

CBMV, LA and PM thank CNPq (The Brazilian National Council for Scientific and Technological Development) for research fellowships. GH was supported by Coordenação de Aperfeiçoamento de Pessoal de Nível Superior (CAPES 88887.671506/2022-00). This study was supported by the Fundação de Amparo à Pesquisa do Estado de São Paulo (FAPESP – 2019/13191-5).

## Conflict of Interest statement

The authors declare that the research was conducted in the absence of any commercial or financial relationships that could be construed as a potential conflict of interest.

## Supplementary data

Supplementary Table 1 - Resistance gene analogs (RGAs) identified in WGCNA module, including RGA family and subgroup classifications.

Supplementary Table 2 - KEGG pathway enrichment results for gene co-expression modules associated with metabolites from Groups 2, 3, and 5.

Supplementary Table 3 - List of differentially expressed genes (DEGs) including log2 fold change values across contrasts and time points.

Supplementary Table 4 - Partial correlation results for gene–metabolite pairs associated with fungal infection and water limitation.

Supplementary Figure 1 - Gene co-expression network construction and module identification using WGCNA. (A) Hierarchical clustering dendrogram of genes based on topological overlap, before and after module merging. Modules were initially defined using a dynamic tree cut method and subsequently merged based on eigengene correlation with a threshold of 0.15. (B) Heatmap and hierarchical clustering of module eigengenes showing the relationships among the 32 gene co-expression modules after merging. Module similarity is indicated by color intensity, with red representing high correlation and blue representing low correlation.

All scripts used in this study are available at the GitHub repository (https://github.com/GustavoHusein/Soybean-integration-Analysis/tree/main/Soybean-integration-Analysis).

## Notes

### Competing Interest Statement

The authors have declared no competing interest.

