## Supplementary figures and images for "Integrative Multi-Omics Analysis Reveals Stress-Specific Molecular Architectures in Soybean under Drought and Rust Infection"

### Supplementary Figure 1

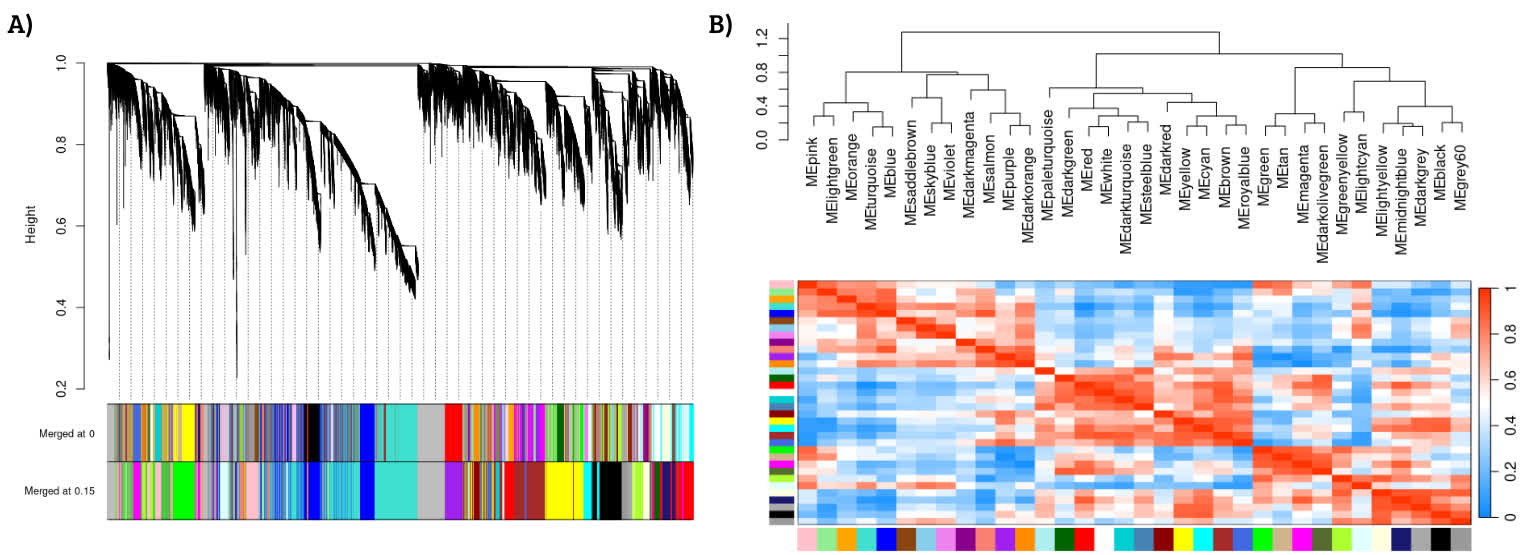
